# A Silicon Diode based Optoelectronic Interface for Bidirectional Neural Modulation

**DOI:** 10.1101/2024.02.27.582240

**Authors:** Xin Fu, Zhengwei Hu, Wenjun Li, Liang Ma, Junyu Chen, Muyang Liu, Jie Liu, Shuhan Hu, Huachun Wang, Yunxiang Huang, Guo Tang, Bozhen Zhang, Xue Cai, Yuqi Wang, Lizhu Li, Jian Ma, Song-Hai Shi, Lan Yin, Hao Zhang, Xiaojian Li, Xing Sheng

**Author notes:** X. F. and Z. H. contributed equally to this work.

## Abstract

The development of advanced neural modulation techniques is crucial to neuroscience research and neuroengineering applications. Recently, optical-based, non-genetic modulation approaches have been actively investigated to remotely interrogate the nervous system with high precision. Here, we show that a thin-film, silicon (Si)-based diode device is capable to bidirectionally regulate in vitro and in vivo neural activities upon adjusted illumination. When exposed to high-power and short-pulsed light, the Si diode generates photothermal effects, evoking neuron depolarization and enhancing intracellular calcium dynamics. Conversely, low-power and long-pulsed light on the Si diode hyperpolarizes neurons and reduces calcium activities. Furthermore, the Si diode film mounted on the brain of living mice can activate or suppress cortical activities under varied irradiation conditions. The presented material and device strategies reveal an innovated optoelectronic interface for precise neural modulations.

**Teaser:** A thin-film, silicon (Si)-based diode device is capable to bidirectionally regulate in vitro and in vivo neural activities.

## INTRODUCTION

Advanced neural modulation techniques are crucial in both research and clinical settings to improve or study functions of nervous systems(*1*). Compared to drug-based biochemical effects, physical cues including electrical, optical, acoustic, magnetic, mechanical and thermal signals provide more controllable and versatile interactions with the biological systems at high spatiotemporal resolutions (*2–7*). In particular, implantable electrical stimulators can be located in targeted neural tissues and modulate cell activities with high precision (*8–10*). Applications of electrical-based neural implants involve the treatment of neurological disorders like Parkinson’s disease(*11, 12*) and the development of prosthetic devices like cochlear implants(*13*). Alternatively, recently developed optogenetic techniques offer an effective means for the specific cell targeting based on the expression of light sensitive ion channels (opsins) (*14, 15*). Nevertheless, optogenetic methods resorts to genetic targeting opsins that hinder their immediate applications in clinics(*16*). In addition, conventional neural stimulators rely on tethered electrodes or fiber optics for electrical or optogenetic interrogations, respectively(*17*).

Moreover, the capability of bidirectional modulation, which is to deterministically excite and inhibit specific neural activities in the same cells or tissues, represents an important focus and is highly desirable for next-generation neural interfaces. In addition to enhancing our understanding of neural function in basic research, bidirectional neural modulation is also potential for innovating therapeutic strategies in the treatment of certain neurological and psychiatric disorders, such as depression, Parkinson’s disease and epilepsy (*18–22*). Traditionally, such a dual modality can be accomplished by injecting inward or outward currents via intracellular or extracellular electrical stimuli, thereby depolarizing or hyperpolarizing the cell membrane potential (*3, 23, 24*). In the aspect of optogenetics, the bidirectional neuronal modulation has been achieved through the co-expression of two spectrally distinct excitatory and inhibitory opsins (for example, ChrimsonR and stGTACR2, or BiPOLES), which activate and silence neural activities by illuminating light at different wavelengths (for example, red and blue), respectively (*25–27*). Alternatively, non-genetic optical approaches, which are based on photocapacitive, photoelectrochemical, photothermal and photoacoustic effects of various materials and devices, have been extensively explored to regulate neural activities (*28–34*). However, these reported concepts only provide a unidirectional modality (either excitation or inhibition). While most studies demonstrate effective neural excitations, non-genetic inhibition capabilities are equally important but have been less investigated. Recently, we have implemented thin-film, monocrystalline silicon (Si) pn junctions as an optoelectronic interface to interrogate the nervous system (*35*). We discovered that these designed Si diodes, under visible or near-infrared illumination, create polarity-dependent optoelectronic signals (photocurrents and photovoltages) in aqueous solution. In particular, optically irradiated n^+^p and p^+^n Si diodes generate positive and negative photovoltages, selectively accumulating anions and cations at the semiconductor/solution interface, respectively. Furthermore, the photoinduced signals can be adjusted by spatially controlled illumination patterns or lithographically defined diode structures, creating dynamic electric field distributions that deterministically activate (p^+^n Si) or inhibit (n^+^p Si) the neural cells or tissues. Nonetheless, the accomplished neural activation or inhibition effect still requires the implementation of two different diode structures (p^+^n or n^+^p Si junctions, respectively) on the nervous system. In other words, non-genetic, optical-based bidirectional modulation in the same cells or tissues, has not yet been realized.

In this work, we present a non-genetic, bidirectional modulation technique that simply utilizes a thin-film Si diode structure to simultaneously activate and inhibit neural activities under optical illumination. By adjusting the optical intensity and pulse width, we discover that such a Si n^+^p diode film can depolarize or hyperpolarize neurons in different conditions. Specifically, high-power and short-pulse irradiations induce transient temperature rise in the Si diode, excite cultured neurons and upregulate intracellular calcium dynamics via photothermal effects. Conversely, low-power and continuous illumination suppresses cell activities and downregulates calcium dynamics through optoelectronic effects. Finally, this Si diode film mounts on the brain of living mice, activating and silencing cortical activities. This bidirectional optical-based neural modulation strategy offers numerous opportunities for next-generation brain machine interfaces.

## RESULTS

### Structural and optoelectronic characteristics of the Si diode film

We employ a thin-film, monocrystalline Si n^+^p diode structure for bidirectional neural modulation (Fig. 1a). The detailed device design and fabrication scheme are provided in Fig. S1 and Methods. Briefly, a thin-film Si n^+^p junction with a thickness of 2 μm is formed by implanting phosphorous ions into p-type silicon-on-insulator (SOI) wafers. The junction depth is measured to be about 0.5 μm, consistent with the simulation result (Fig. S2). Selective wet-etching releases the lithographically patterned Si film from the original wafer, and transfer printing integrates the Si film onto heterogeneous (glass or flexible) substrates (*35*). For improved optoelectronic responses in the aqueous environment, gold nanoparticles (AuNPs) are deposited on the p-side of the junction, which is in direct contact with the neural cells or tissues (*36*).

**Fig. 1.**
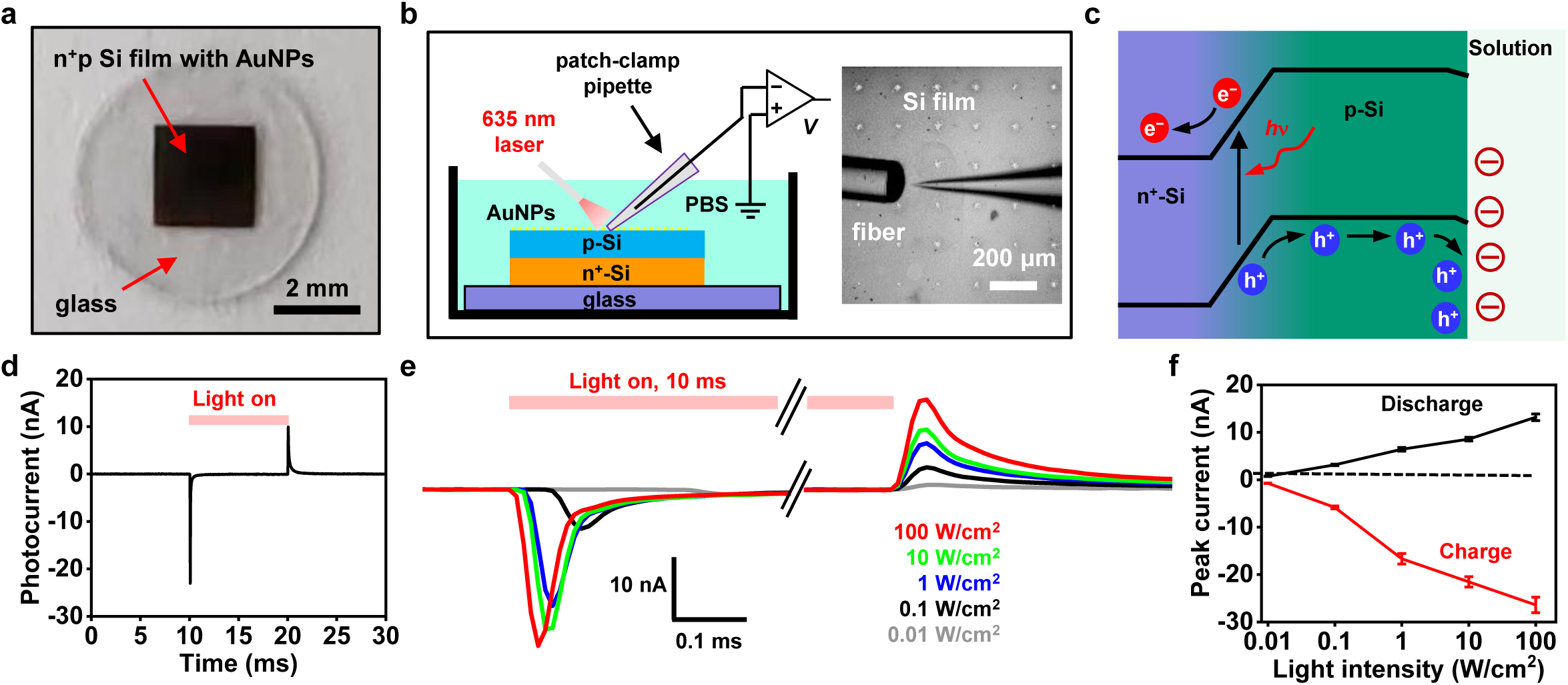
In vitro optoelectronic response of an n^+^p Si diode film with decorated gold nanoparticles (AuNPs). (a) Picture of an n^+^p Si film with AuNPs (thickness ∼2 µm, lateral dimension ∼2 mm) transferred on glass. (b) (left) Scheme and (right) microscopic image of the patch-clamp setup to measure the photocurrent of the Si film. (c) Schematic illustration of photogenerated carriers (electrons e^-^ and holes h^+^) within the Si junction and the ion accumulation at the Si/solution interface. (d) Photocurrent response for the Si film. The illumination condition is: intensity 10 W/cm^2^, pulse duration 10 ms. (e) Exploded view of charge and discharge peak currents for the Si film under illumination with different intensities (0.01, 0.1, 1, 10, 100 W/cm^2^). (f) Peak currents of charge and discharge as a function of light intensities. Data are presented as mean ± s.e.m..

We use a customized patch-clamp setup to characterize the in vitro optoelectronic response of the AuNP-decorated Si diode film in the phosphate-buffered saline (PBS) solution (Fig. 1b). Optical illumination (at a wavelength of 635 nm) incident on the Si film creates photogenerated carriers (electrons and holes) and establishes an electrical field, which causes the ion accumulation at the Si/solution interface (Fig. 1c) and eventually leads to photocapacitive currents (Fig. 1d). Specifically, light-on and light-off states elicit charge and discharge currents, respectively, and both of them increase monotonically with the light intensity up to ∼100 W/cm^2^ (Figs. 1e and 1f). Furthermore, photocapacitive currents produced by the Si diode film with AuNPs are substantially larger than those for the device without AuNPs (Fig. S3), and no photo response is observed when similar irradiation is imposed on glass (Fig. S4).

### Photothermal characteristics of the Si diode film

The same patch-clamp setup in Fig. 1b is also capable to capture the photothermal effect of the Si film under illumination, based on the temperature-dependent resistance change of the pipette electrode (*37*). Described in Methods and Fig. S5, the patch-clamp probe detects the dynamic temperature change (at the millisecond scale) at the semiconductor/solution interface with a high positioning accuracy, which is difficult to accomplish with other thermal characterization methods based on commercial infrared imagers or thermocouples. Under illumination with an intensity of 100 W/cm^2^ and varied pulse widths (1, 2, 5, and 10 ms), the maximum temperature rises range from ∼3 K to ∼15 K (Figs. 2a and 2c). When we fix the pulse width to 10 ms and change the intensity (10, 50, 75, and 100 W/cm^2^), the maximum temperature rises range from ∼2 K to ∼15 K (Figs. 2b and 2d). These measured photothermal results are also in accordance with those predicted by numerical simulations based on the finite-element analysis (Figs. 2e and 2f). Fig. S6 presents the photothermal characteristics for a Si diode film without AuNPs, which exhibits higher temperature increases than the sample with AuNPs under similar illumination conditions. This difference may be caused by the different optical reflectance on Si surfaces with and without AuNPs. Under continuous irradiance, the thermal effects should be mitigated in order to avoid irreversible tissue damage induced by prolonged overheating (*38, 39*). For a 5-s pulse duration and varied intensity from 2.5 to 12.5 W/cm^2^, the temperature change is between ∼1 K and ∼5 K, revealed by both experimental and simulation results (Figs. 2g and 2h). The agreement between measured and simulated data indicates that the numerical calculation can provide a quantitative prediction on the photothermal effects of the Si film immersed in aqueous solutions. Fig. 2i summarizes the calculated maximum temperature rise as a function of pulse duration (from 0.1 ms to 10 s) and light intensity (from 0.1 to 100 W/cm^2^). Such a map can offer a guideline to predict the thermal impact on the biological cells or tissues attached to the Si film.

**Fig. 2.**
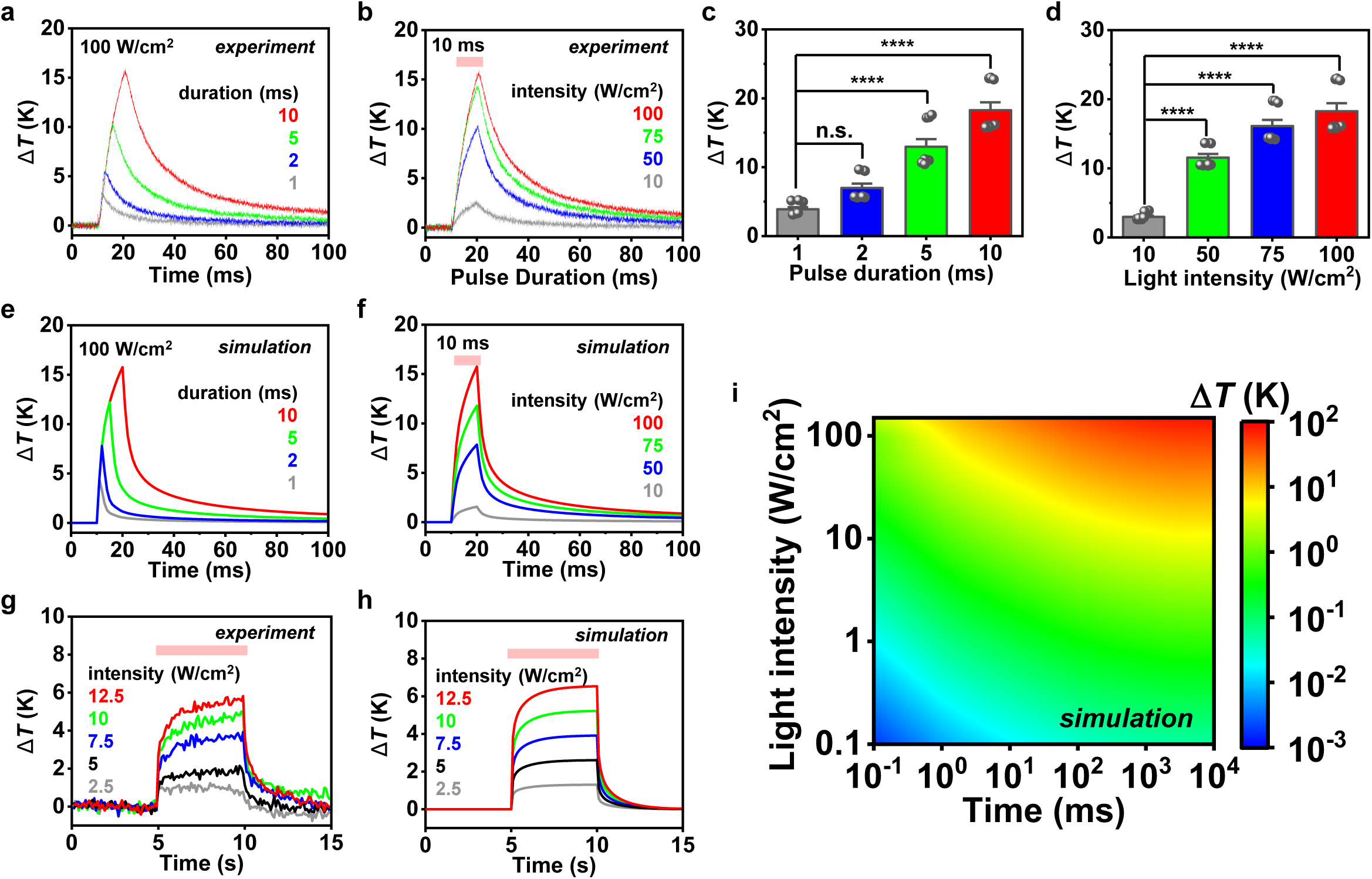
In vitro photothermal response of the Si diode film in the PBS solution. (a, b) Measured transient temperature increase (Δ*T*) of a Si film illuminated by a pulsed laser at 635 nm, (a) with an intensity of 100 W/cm^2^ and different pulse durations (1, 2, 5, 10 ms), (b) with a pulse duration of 10 ms and different intensities (10, 50, 75, 100 W/cm^2^). (c, d) Statistical data of the maximum temperature rise related to conditions in (a) and (b) (*n* = 3 devices, 3 trials for each). (e, f) Simulated transient temperature response of a Si film corresponding to experimental conditions in (a) and (b). (g) Measured and (h) Simulated dynamic temperature increase of a Si film illuminated by a laser at 635 nm with a pulse duration of 5 s and different intensities (2.5, 5, 7.5, 10, 12.5 W/cm^2^). (i) Simulated map of maximum temperature rise as a function of pulse duration and light intensity. Data are presented as mean ± s.e.m and analyzed by one-way RM ANOVA. \**p* < 0.05, \*\**p* < 0.01, \*\*\**p* < 0.001, \*\*\*\**p* < 0.0001, n.s., not significant.

### Bidirectional modulations of electrophysiological activities for cultured neurons

To evaluate the influence of illumination on neural activities, we culture rat dorsal root ganglion (DRG) neurons on the AuNP-decorated n^+^p Si diode film (Figs. 3a). Before cell culture, the device is treated with oxygen plasma and coated with poly-L-lysine (PLL) and laminin, which create a hydrophilic surface and promote cell adhesion (Fig. S7). This pretreated device sample exhibit a desirable biocompatibility with cultured DRG neurons, which have viability comparable to those on the standard glass substrate (Fig. S8).

**Fig. 3.**
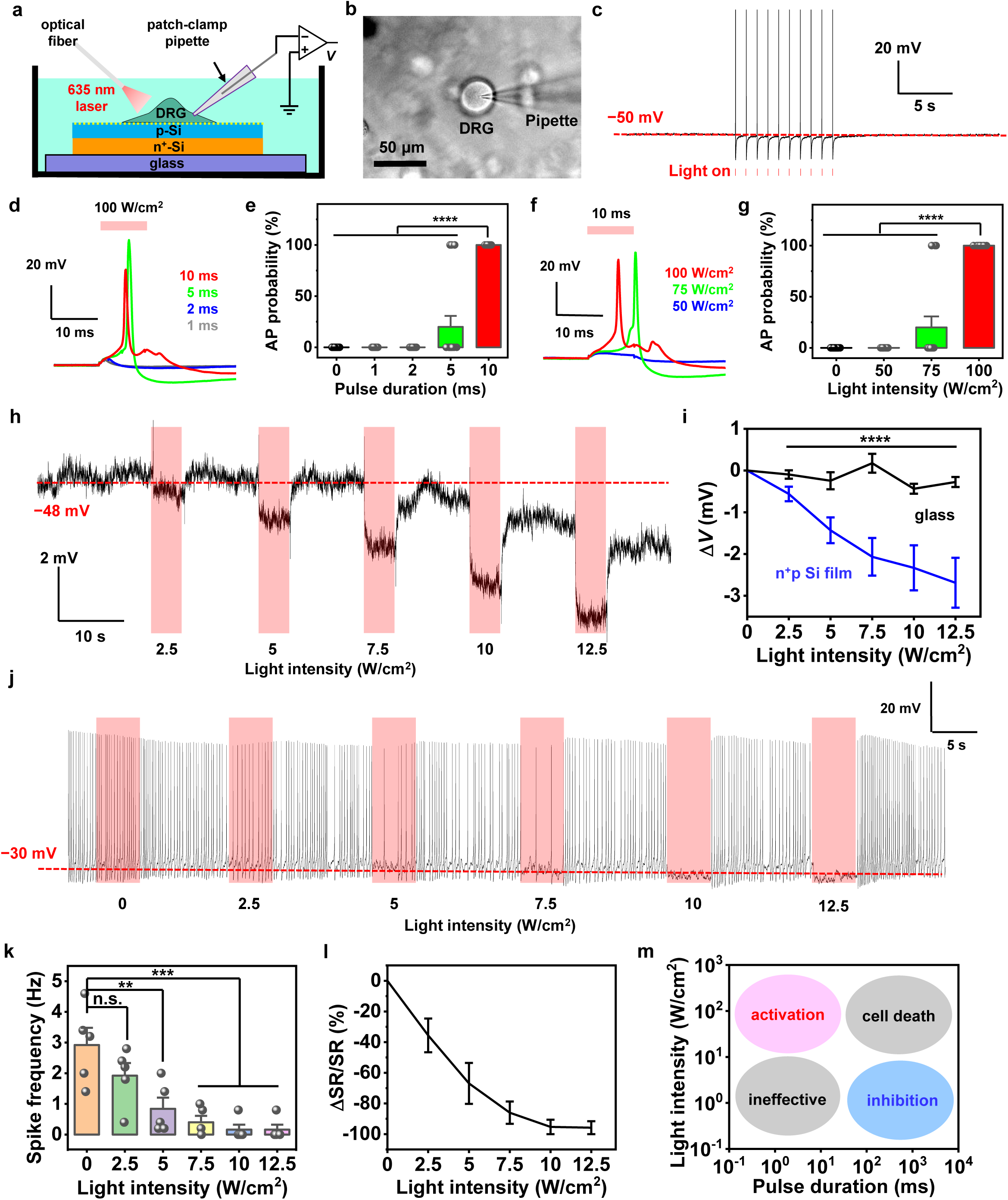
Bidirectional activation and inhibition of electrophysiological activities of cultured dorsal root ganglion (DRG) neurons with the Si diode film. (a) Schematic illustration of the patch-clamp setup to record electrophysiological signals of DRG neurons cultured on the Si film under light stimulation (with a 635 nm laser, ∼0.5 mm spot size). (b) Microscopic image of a DRG cell recorded by a pipette electrode. (c) Representative trace of action potentials (APs) activated by pulsed light (intensity 100 W/cm^2^, duration 5 ms, frequency 1 Hz). (d) Dynamic changes of membrane potentials and (e) the probability of APs in response to pulsed light (intensity 100 W/cm^2^, durations 1, 2, 5, 10 ms) (*n* = 5 neurons, 3 trials for each neuron). (f) Dynamic changes of membrane potentials and (g) the probability of APs in response to pulsed light (duration 10 ms, intensity 50, 75, 100 W/cm^2^) (*n* = 5 neurons, 3 trials for each). (h) Representative trace showing hyperpolarization of the DRG cell modulated by continuous illumination (duration 5 s, intensities 2.5, 5, 7.5, 10, 12.5 W/cm^2^). (i) Statistical results showing the maximum decrease of hyperpolarized membrane voltages under light stimulation with different intensities (Si film: *n* = 4 neurons, 9 trials in total; glass substrate: *n* = 4 neurons, 6 trials in total). (j) Representative trace showing that APs of a DRG neuron are suppressed by the Si diode film with different light intensities. The cell is initially excited by injecting currents with a pipette electrode. (k) Statistics of spike frequency before and during the stimulation (5 s) with different intensities (*n* = 4 neurons, 5 trails in total). (l) Statistics of spike rate change (△*SR*/*SR*) with different light intensities. △*SR*/*SR* is calculated by: [*SF*_(during)_ − *SF*_(before)_]/*SF*_(before)_, where *SF*_(during)_ and *SF*_(before)_ represent the averaged spike frequencies (*SF*) during and before the stimulation, respectively (*n* = 4 neurons, 5 trails in total). (m) Schematic illustration showing different regulation effects of the Si film on the cell behaviors at different pulse durations and light intensities. Data are presented as mean ± s.e.m. and analyzed by one-way RM ANOVA. \**p* < 0.05, \*\**p* < 0.01, \*\*\**p* < 0.001, \*\*\*\**p* < 0.0001, n.s., not significant.

We study the photoinduced cell activities using the whole-cell patch recording (Fig. 3b). Optical irradiance with different intensities and pulse widths is incident on DRG neurons on the Si film. We first find that high-intensity, short-pulsed illumination (for example, 100 W/cm^2^, 5 ms) can activate DRGs and induce action potentials (APs, or spikes) that follow each pulse (Fig. 3c). When we reduce the light intensity or the pulse width, cell depolarization effects are diminished. If the membrane potential does not reach the threshold voltage, the induced cell depolarization cannot result in a complete spike (Fig. 3d-3g). Illumination with a pulse width of 10 ms and an intensity of 100 W/cm^2^ can elicit spikes with almost 100% possibility, while reducing the pulse width to 5 ms or the light intensity to 75 W/cm^2^ only induces APs in some cells. The observed cell activation can be ascribed to the rapid temperature increase associated with the light pulse incident on the Si film (demonstrated in Fig. 2), and similar photothermal effects are explored for Si nanostructures and other light absorptive materials (*40–48*). Photo responses of these DRG neurons are distinct from those of cells cultured on the Si diode with the opposite polarity (p^+^n film), which generates a more slowly enhanced membrane potential and leads to a greatly delayed (by a few seconds) AP under much lower intensities (*35*). Another interesting finding is that optically induced APs for neurons on the Si film show much lower peak potentials (∼10 mV) comparing to the typical APs initiated by conventional electrical-based intracellular stimulus (Fig. S9). This result may be attributed to the negative photovoltages generated by the n^+^p Si film (*35*), which compensates a portion of the elevated membrane potential. It is also noted that higher illumination intensity (> 100 W/cm^2^) can have phototoxicity, eventually causing irreversible cell damage and death (Fig. S10).

The same AuNP-decorated n^+^p Si diode film also produces neural inhibitive function, under low-intensity, prolonged illumination. Shown in Figs. 3h and 3i, continuous illumination (5 s) on the Si diode film reduces membrane potentials of DRG neurons, by values from 0.5 mV to 3 mV with light intensities varying from 2.5 W/cm^2^ to 12.5 W/cm^2^. The cell hyperpolarization could be owing to the photocapactive effect of the n^+^p Si diode (*35*). Furthermore, such photoinduced hyperpolarization can suppress electrically induced APs in DRG neurons (Fig. 3j). When the irradiance increases up to 12.5 W/cm^2^, the spike frequency gradually decreased from 3 Hz to 0, and spikes are fully inhibited at 10 or 12.5 W/cm^2^ (Figs. 3k and 3l).

The above results suggest that photothermal and photocapacitive responses of the AuNP-decorated n^+^p Si diode film co-exist and exert different influences on the neural activities, depending on the illumination intensity and the pulse duration. Fig. 3m provides a qualitative map to illustrate these diverse outcomes. High-power and short-pulsed light excites the DRG neurons, mostly due to the transient temperature rise of the Si film. On the contrary, low-power and long-pulsed light dominantly create optoelectronic signals that inhibit the cells. On the other hand, low-power and short-pulsed light simply does not alter the cell function, and high-power and prolonged irradiance inevitably causes cell dysfunction and death. As a comparison, similar irradiations imposed on DRG neurons cultured on transparent and photo-insensitive glass substrate do not induce any depolarization or hyperpolarization effects under similar illuminations (Fig. S11).

### Bidirectional modulations of calcium dynamics for cultured neurons

Calcium (Ca^2+^) signals directly correlate with the electrophysiological activities of neural cells. In Fig. 4, we capture Ca^2+^ dynamics of DRG neurons cultured on the AuNP-decorated n^+^p Si diode film in response to light. A confocal microscope records the intracellular Ca^2+^ fluorescence of neurons loaded with the Ca^2+^ indicator (Calbryte-520). In this experiment, we investigate the behavior of a same group of DRG neurons on the Si film upon high-power (100 W/cm^2^), short-pulsed (10 ms) (Fig. 4a) or low-power (1 W/cm^2^) and long-pulsed (20 s) illuminations (Fig. 4h). Corresponding images and videos are presented in Fig. 4b and 4g, Movie S1 and S2, respectively. The high-power and short-pulsed light triggers increased Ca^2+^ fluorescence of multiple DRG neurons, by a factor of about 5 (Figs. 4c-4f). Conversely, Ca^2+^ activities of these cells are suppressed by low-power and long-pulsed irradiation, by about 10% (Figs. 4h-4j). These activated and inhibited Ca^2+^ dynamics are in accordance with the electrical activities of neurons responding to the optical modulations, validating the bidirectional functionality of the Si diode film.

**Fig. 4.**
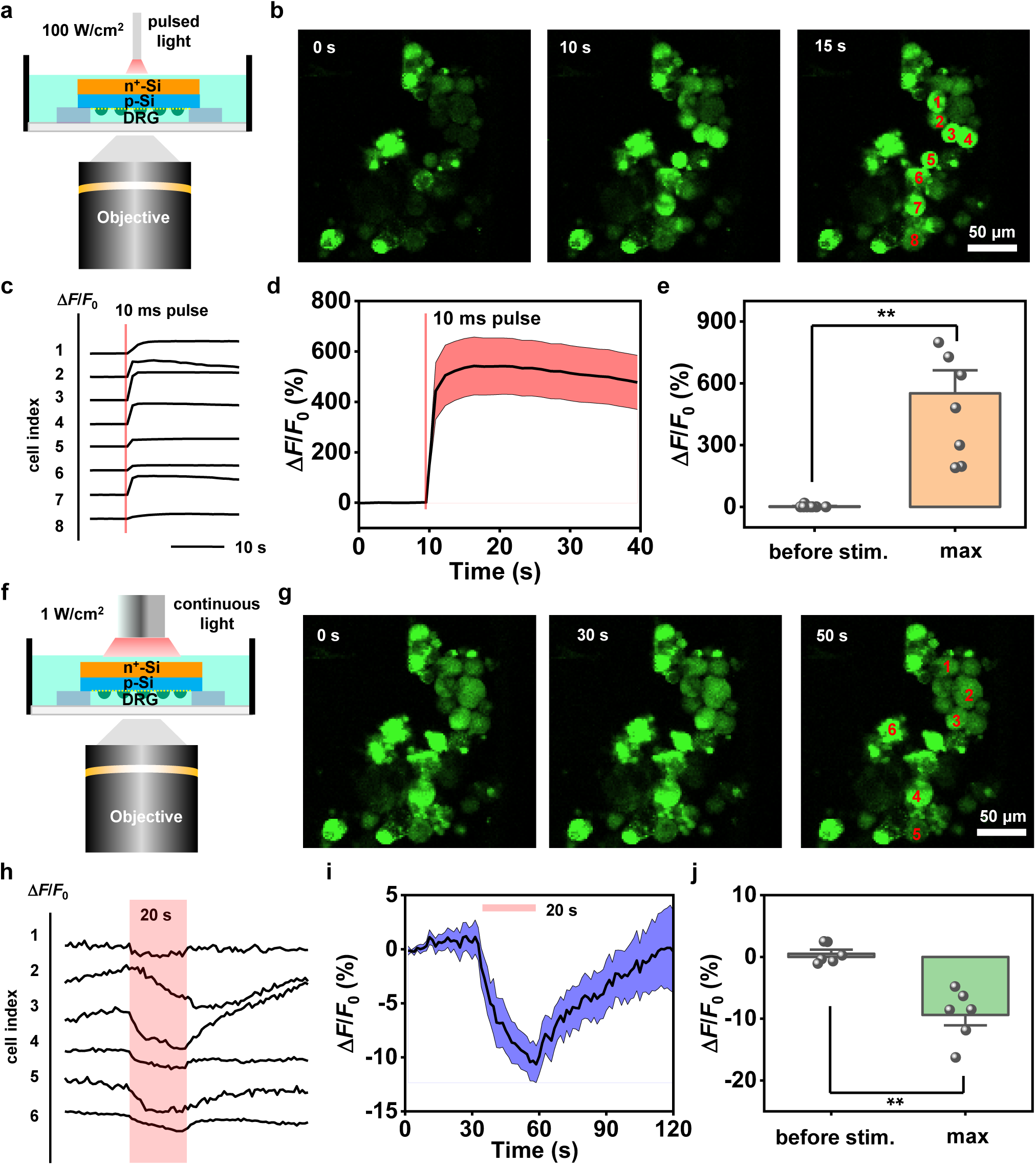
Bidirectional modulation of calcium (Ca^2+^) dynamics of DRG neurons with the Si diode film. (a) Illustration of the microscopic setup to image the Ca^2+^ fluorescence of DRG neurons, which are activated by illuminating a pulsed light on the Si film (light intensity 100 W/cm^2^, duration 10 ms). (b) Fluorescent images of DRG neurons cultured on the Si film, at different time frames before and after optical stimulation. (c) Transient Ca^2+^ signals (Δ*F*/*F*_0_) of multiple DRG neurons circle-marked in (b). (d) Average Ca^2+^ signals. The solid lines and shaded areas indicate the mean and s.e.m., respectively. (e) Comparison of maximum Ca^2+^ signals (Δ*F*/*F*_0_) before and after pulsed stimulation for all marked neurons (*n* = 6 neurons). (f) Illustration of the microscopic setup to image the Ca^2+^ fluorescence of DRG neurons, which are inhibited by continuous illumination on the Si film (light intensity 1 W/cm^2^, duration 20 s). (g) Fluorescent images of DRG neurons cultured on the Si film, at different time frames before and during optical stimulation. (h) Transient Ca^2+^ signals (Δ*F*/*F*_0_) of multiple DRG neurons circle-marked in (g). (i) Average Ca^2+^ signals. The solid lines and shaded areas indicate the mean and s.e.m., respectively. (j) Comparison of maximum Ca^2+^ signals (Δ*F*/*F*_0_) before and during stimulation for all marked neurons (*n* = 8 neurons). All data are presented as mean ± s.e.m. and analyzed by unpaired *t*-test. \**p* < 0.05, \*\**p* < 0.01, \*\*\**p* < 0.001, \*\*\*\**p* < 0.0001, n.s., not significant.

### Bidirectional modulations of cortical neural activities in vivo

In addition to the demonstration on cultured cell in vitro, we also mount the Si diode film on the cortical region of rats and perform optical modulations in vivo (Fig. 5). A released n^+^p Si diode film (2 × 2 mm^2^) is bonded on a flexible polyethylene terephthalate (PET) substrate (Fig. 5a) and subsequently attached on the somatosensory cortex of a living rat, with the AuNP-decorated p-side in contact with the tissue. A 32-channel Si-based microelectrode array is inserted into the brain tissue underneath the Si film and captures extracellular activities, while a laser beam is delivered on the Si film (Figs. 5b and 5c). Fig. 5d presents a representative unit with sorted neural spikes, indicating the quality of the recording. First, we implement high-power, short-pulsed laser (100 W/cm^2^, 10 ms) and record corresponding cortical activities. In this test, multiple trials are collected and sorted, and the generated raster plot and heatmap of cellular activities evidence significantly increased firing spikes after the irradiation (Figs. 5e and 5f). Within the pulse period (10 ms), we observe both activated and silenced units, which could be attributed to the existence of different cell types of cortical neurons and their dissimilar responses in the complex network. Nevertheless, most neurons exhibit clearly enhanced firing rates after illumination, compared to their basal activities prior to stimulation (Fig. 5g). Next, we observe that cortical activities can be inhibited when applying a low-power, long-pulsed illumination (10 W/cm^2^, 1 s) on the Si diode film (Figs. 5h-5j). Specifically, the irradiation is imposed when the animal is experiencing high-level spontaneous activities (averaged firing rate ∼1 Hz). Both the raster plot and the heatmap collected among multiple units indicate greatly suppressed extracellular activities over the course of the modulation (Figs. 5h-5j). As for the control group, similar illumination conditions directly applied on the rat cortex (without Si film) do not evoke any excitation or inhibition responses on in vivo neural activities (Figs. 5g and 5j).

**Fig. 5.**
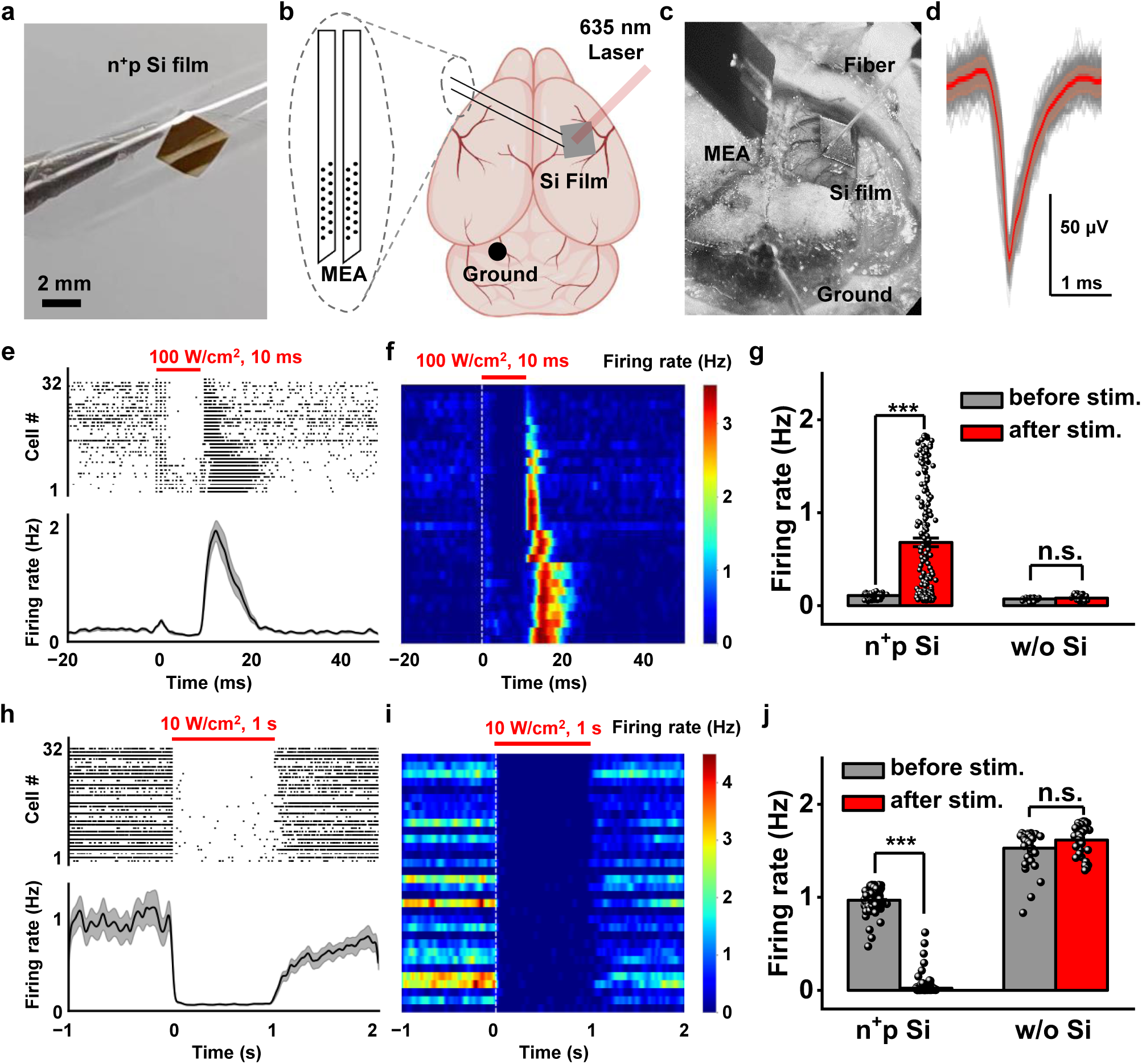
Bidirectional activation and inhibition of in vivo cortical activities with the Si diode film. (a) Photograph of the Si film attached on a flexible polyethylene terephthalate (PET) film. (b) Schematic illustration and (c) Photograph of the experiment setup. The Si film is mounted on the rat cortex and illuminated by light, while a multichannel electrode array is inserted beneath the Si film to record the cortical electrophysiological signals. (d) Representative aligned waveforms recorded from a neuron. Each black line depicts one spike waveform and the red line depicts the averaged waveform. (e) Spike raster plots collected from 32 recording channels (top) and averaged neuron firing rates (bottom) in response to a 10-ms pulsed stimulation. (f) Heatmap of 32 channels. Statistics of firing rates before and after stimulations for cells on Si and without Si (before: [−20, 0] ms, after: [10, 30] ms. Si film: *n* = 32 neurons from 5 rats, Without Si: *n* = 6 neurons from 3 rats). Spike raster plots collected from 32 recording channels (top) and averaged neuron firing rates (bottom) in response to a 1-second stimulation. (i) Heatmap of 32 channels. (j) Statistics of firing rates before and during stimulations for cells on Si and without Si (before: [−1, 0] s, after: [0, 1] s, Si film: *n* = 32 neurons from 5 rats, Without Si : *n* = 12 neurons from 3 rats). All data are presented as mean ± s.e.m., and analyzed by Mann-Whitney U test. \**p* < 0.05, \*\**p* < 0.01, \*\*\**p* < 0.001, \*\*\*\**p* < 0.0001, n.s., not significant.

## DISCUSSION

To summarize, we demonstrate that the AuNP-decorated n^+^p Si diode film enables the simultaneous optical-based, non-genetic neural activation and inhibition, in both in vitro and in vivo experiments. In particular, we find that both photothermal and photocapacitive effects occur at the interface between the semiconductor and the neurons, and they evoke cell depolarization or hyperpolarization depending on different illumination conditions. Further studies are required to fully understand the fundamental mechanism of such a bidirectional modulation method, which may be associated with the co-existence of temperature- and voltage-sensitive ion channels in neurons (*49*), as well as their complicated interactions with the Si diode upon illumination. In the future, geometrical structures of the Si film and illumination patterns can be more precisely tuned, to interrogate individual cells or neural networks with higher spatial and temporal resolutions. Application scenarios of this non-genetic modulation strategy include both in vitro and in vivo fields, ranging from cultured cells and organoids to peripheral and central nervous systems. The bidirectional regulation would be promising for use in areas like drug screening (*50*), retinal prosthesis (*10, 51*), tissue regeneration (*52*) and motorsensory restoration (*53*). Collectively, our work here not only sheds light on the unusual interactions between semiconductor and biological systems, but also provides possible insight on next-generation brain-machine interfaces.

## MATERIALS AND METHODS

### Material and device preparation

The n^+^p Si junction was made of an SOI wafer (device layer, p-type, (100), 1-10 Ω cm, ∼2 μm; buried oxide layer, 1 μm; handle layer, p-type, (100), 1-10 Ω cm, 525 μm) followed by implantation of phosphorus (P) (dose 4 × 10^14^ ions cm^−2^, energy 75 keV). After acid cleaning, the implanted wafers were annealed for 30 mins at 950 °C for dopant activation.

Patterns of doped Si films were lithographically defined by reactive ion etching. The SOI wafer was cleaned in acetone for 10 min with ultrasonic cleaning, then in H_2_O_2_ : NH_4_OH : H_2_O =1:1:5 (10 min, 80 °C). SOI wafer was etched in hydrofluoric acid (40% HF, ACS grade, Aladdin) for 3 min. Then SPR 220 was used to make an anchor. The SOI was dipped in the hydrofluoric acid (40% HF, ACS grade, Aladdin) over night to remove the buried oxide layer and release the 2-µm-thick Si films (size 2 ×2 mm^2^).

Polydimethylsiloxane (Dow Corning Sylgard 184 kit, 1:10 weight ratio) stamps and thermal release tapes (No.3198, Semiconductor Equipment Corp.) were applied to transfer the Si films onto target substrates. An epoxy (SU8-3005, 5 µm thick) was spinned on substrates to form an adhesive layer. After transferring Si films, samples were baked at 180 °C for 2 hours to detoxify the SU8 film (*54, 55*). For patch clamp, Si films were transferred on glass coverslip (thickness ∼150 µm, diameter ∼ 6 mm). For calcium imaging, Si films were transferred on glass coverslip (thickness ∼150 µm, diameter ∼ 6 mm) with a hole (diameter 1.5 mm). For optical modulations in mice’s cerebral cortex, Si films were transferred on a PET film (thickness ∼20 µm).

The device was coated with gold nanoparticles, by dipping into 0.5 mM HAuCl_4_ (No.G4022, Sigma-Aldrich) in 1% HF (GR, 40%, Aladdin) for 3 min. Finally, the device was treated with oxygen plasma for 90 seconds to make the surface more hydrophilic and promote cell adhesion (*56, 57*). Before cell culture, the device was coated with 0.01% poly-l-lysine (No.P8920, Sigma-Aldrich) overnight, then coated with 1∼2 μg cm^−2^ laminin (No. L2020, Sigma-Aldrich) for more than 4 hours. The water contact angle on the sample surface was measured by a general-purpose video optical contact angle tensiometer (LSA60, Germany LAUDA Scientific)

### Optoelectronic characterizations

A standard patch-clamp setup was used to measure the photon response of the Si films. Micropipettes were pulled from glass capillaries using a micropipette puller (P-97, Sutter Instruments) for a final resistance of ∼2 MΩ in 1× phosphate buffered saline (1× PBS) solution (Solarbio, P1020). During the photothermal characterization, the dish (Corning, 430165) was filled with 4 mL of 1× PBS solution and a Ag/AgCl wire was placed in the chamber. Current-clamp protocols were performed with an Axopatch 700B amplifier controlled by pClamp software (Molecular Devices). Glass pipettes (∼1–2 MΩ) filled with 1× PBS solution is positioned about 1 μm above the Si surface. Light stimulations were generated by a 635 nm laser (MDL-D-635 nm, Changchun New Industries Optoelectronics Tech. Co., Ltd.), with controlled power, frequency and duration. A laser beam (spot size ∼500 μm) was delivered on the Si film through an optical fiber (diameter 200 μm). The light pulses were controlled by patch clamp logic signals. The light intensity from the fiber tip was measured by using a power meter (LP10, Sanwa). The transient photoresponse measurements were taken by voltage-clamp recording (filtered at 10 kHz and sampled at 100 kHz), with Si films fully immersed in PBS solution.

### Photothermal characterizations

Photothermal characterization of Si film was performed using a previously described micropipette method in a voltage-clamp mode (*37*). To calibrate the resistance of the micropipette as a function of temperature, 4 mL of 1×PBS solution was heated to 60 °C and added into a 35 mm dish. The temperature of the solution was continuously measured by a T type thermal couple using a temperature logger (USB-TC01, National Instruments Co.) to establish a calibration curve, the micropipette’s resistance was recorded until a bath temperature of 20 °C was reached (Fig. S5a and S5b). The dependence between the micropipette’s resistance *R* and temperature *T* follows the Arrhenius equation (*58*):

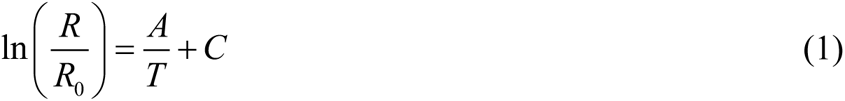

According to Ohm’s law:

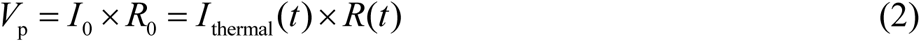

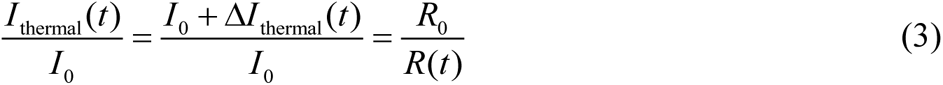

where *V*_p_, *I*_0_, *R*_0_, *I*_thermal_(*t*), *R*(*t*), are the holding voltage (14 mV), the injected current (7.5 nA), the micropipette resistance at 20 °C, the thermal current, the measured micropipette resistance, respectively. The transient photocurrent Δ*I*_photo_(*t*) was collected with without bias current (Fig. S5c). Δ*I*_thermal_(*t*) was obtained by subtracting Δ*I*_photo_(*t*) and *I*_0_ from the overall photocurrent, and *R*(*t*) was determined based on Eq.(3) (Fig. S5d and S5e). Finally, the temperature *T* was determined by the fitted ln*R* − 1/*T* curve (Fig. S5f).

### Photothermal simulation

A finite-element analytical model (COMSOL Multiphysics) is used to calculate transient temperature rise of silicon film in response to irradiation. The materials and corresponding parameters (density, thermal conductivity and heat capacity) used in the model include: silicon (2.3 g cm^−3^, 150 W m^−1^ K^−1^, 0.71 J g^−1^ K^−1^), glass (2.2 g cm^−3^, 1.4 W m^−1^ K^−1^, 0.73 J g^−1^ K^−1^), water (1.0 g cm^−3^, 0.61 W m^−1^ K^−1^, 4.2 J g^−1^ K^−1^) and SU-8 (1.2 g cm^−3^, 0.2 W m^−1^ K^−1^, 1.5 J g^−1^ K^−1^). The light absorption of the Si film is set to 60% and the diameter of light spot is 0.5 mm. The results of transient temperature rise of silicon film under the same total power irradiation for different times and under the same irradiation time but different total power are simulated. Subsequently, the temperature rise model of the silicon film modified with AuNPs is constructed. Considering that the light reflection of the scale surface is enhanced after the addition of AuNPs, the reflectivity is set at 25%, and the relationship between temperature change and time is obtained as with the former.

### Culture of DRG neurons

DRGs of male rats (4 weeks) were extracted and transferred in ice-cold of DMEM/F12 (No.11330-032, gibco). Tissues were cut into small pieces and treated with an enzyme solution containing 5 mg ml^−1^ dispase (No.ST2339, beyotime) and 1 mg ml^−1^ collagenase (No.ST2294, beyotime) at 37 °C for 1 h. After trituration with 1 ml pipette for 30 times until the liquid becoming turbid. After centrifugation at 250 g for 5 min, cells were washed in 15% (w/v) bovine serum albumin (SRE0096-10G, Sigma) and resuspended in 2 ml of DMEM/F12 containing 10% fetal bovine serum, and 1% penicillin–streptomycin (15140-122, gibco). One tenth of the total DRG cells was planted on the device in each well of the 96 wells plate. The DRG cells was cultured in an incubator at 37 °C for 1 day (*59*).

### Viability Assay

For the assay, the samples were labeled with LIVE/DEAD™ Cell Imaging Kit (Invitrogen). The live cells were labeled with Calcein AM (green); the dead cells were labeled with EthD-1(red); After adding the dyes, the samples were incubated for 15 min at 37 °C with 5% CO_2_. The samples were washed with HBSS for three times. The viability (%) was calculated as total no. of live cells / total No. of cells ×100.

### In vitro electrophysiological recording

DRG neurons were perfused with artificial cerebrospinal fluid (ACSF) containing the following (in mM): 126 NaCl, 3 KCl, 1 MgCl_2_, 2.5 CaCl_2_, 10 glucose, 26 NaHCO_3_ and 1.2 NaH_2_PO_4_ at room temperature. Recording pipettes (6–7 MΩ) were pulled with a micropipette puller (P1000, Sutter Instrument). For whole-cell recordings, pipettes were filled with internal solution that contained the following (in mM): 120 K-gluconate, 10 HEPES, 0.2 EGTA, 2 KCl, 4 Na_2_ATP, 0.4 Na_2_GTP, 2 MgCl_2_ and 10 Na_2_phosphocreatine (pH 7.2–7.4). Voltage-and current-clamp recordings were performed with a computer-controlled amplifier (MultiClamp 700B, Molecular Devices). Recorded traces were low-pass filtered at 3 kHz and sampled at 10 kHz (DigiData 1440, Molecular Devices). Collected data were analysed using Clampfit 10 software (Molecular Devices).

### Calcium imaging

Cells were loaded with Calbryte 520 (Screen Quest™ Calbryte-520™ Probenecid-Free and Wash-Free Calcium Assay Kit, AAT-A36317) at 37 °C for 1 h. Then the device was removed from the 96 wells and placed on the confocal dish (diameter 35 mm) with the cells above the glass for 150 μm. Almost 200 μl HBSS was added to the confocal dish. Ca^2+^ fluorescent signals were imaged with a 20×objective on an inverted confocal microscope (Nikon A1R HD25). Fluorescence images were analysed via ImageJ. Normalized fluorescence changes were calculated as Δ*F*/*F*_0_ = (*F* − *F*_0_)/*F*_0_, where *F*_0_ is the baseline intensity.

### Stimulation and recording cortical activities in vivo

Mature adult Sprague Dawley rats (weights on around 200 g) were used in this study. Rats were maintained at room temperature (24 °C) with proper humidity and on a 12-hour light/dark cycle and fed at libitum. All animal protocols used were in accordance Tsinghua University and the Shenzhen Institute of Advanced Technology, and were approved by their Institutional Animal Care and Use Committee.

Rats were firstly anesthetized with 2% isoflurane and maintained with pentobarbital sodium (1% kg^−1^). After mounted on the stereotaxic apparatus (RWD, China) using ear bars, eye ointment was covered on the animals to protect from desiccation. The exposed skull was under a craniotomy to tile the film. Stainless steel screw soldered the silvery wire were used as ground on the cerebellum. The silicon electrode was aslant (∼25°–30°) inserted along the near edge of the Si film, and the optic fiber was positioned vertically above small portion of the Si film.

Neural activity in rats was recorded using a self-designed system and silicon microelectrodes (Lotus, American). We employed traditional approaches for spike detection, dividing the entire procedure into three steps: noise removal, thresholding detection, and average template matching. Firstly, band pass filtering was used with cutoff frequency ranging from 250 to 7000 Hz. The desired spike trains are primarily contained in the remained signal. Due to the laser artifacts when recording, the technique of Fast Fourier Transform was specifically involved to remove of such laser-induced noise. After denoising step, we needed a simple but preliminary detection for the potential spikes. We assumed that the peak of existence putative spike often exceeds −60 mV, and the consecutive spikes probably not occurs in an extremely short period (inter-spike-interval is larger than 1 ms). Finally, for the purpose of improvement in the quality of detected spikes, we further implemented an average template matching algorithm. Each spike identified during the thresholding process was measured for its similarity to the averaged templates. The templates for each channel were obtained from visually better spikes that are identified in the previous step. Noisy channels and those with low firing through the recording were discarded. Spikes were grouped according the repetitive trials. Neural firing rate were smoothed using gaussian filter for visualization purposes only. All the analysis were conducted using Python 3.10.

## Supplementary Materials

**Figure S1.**
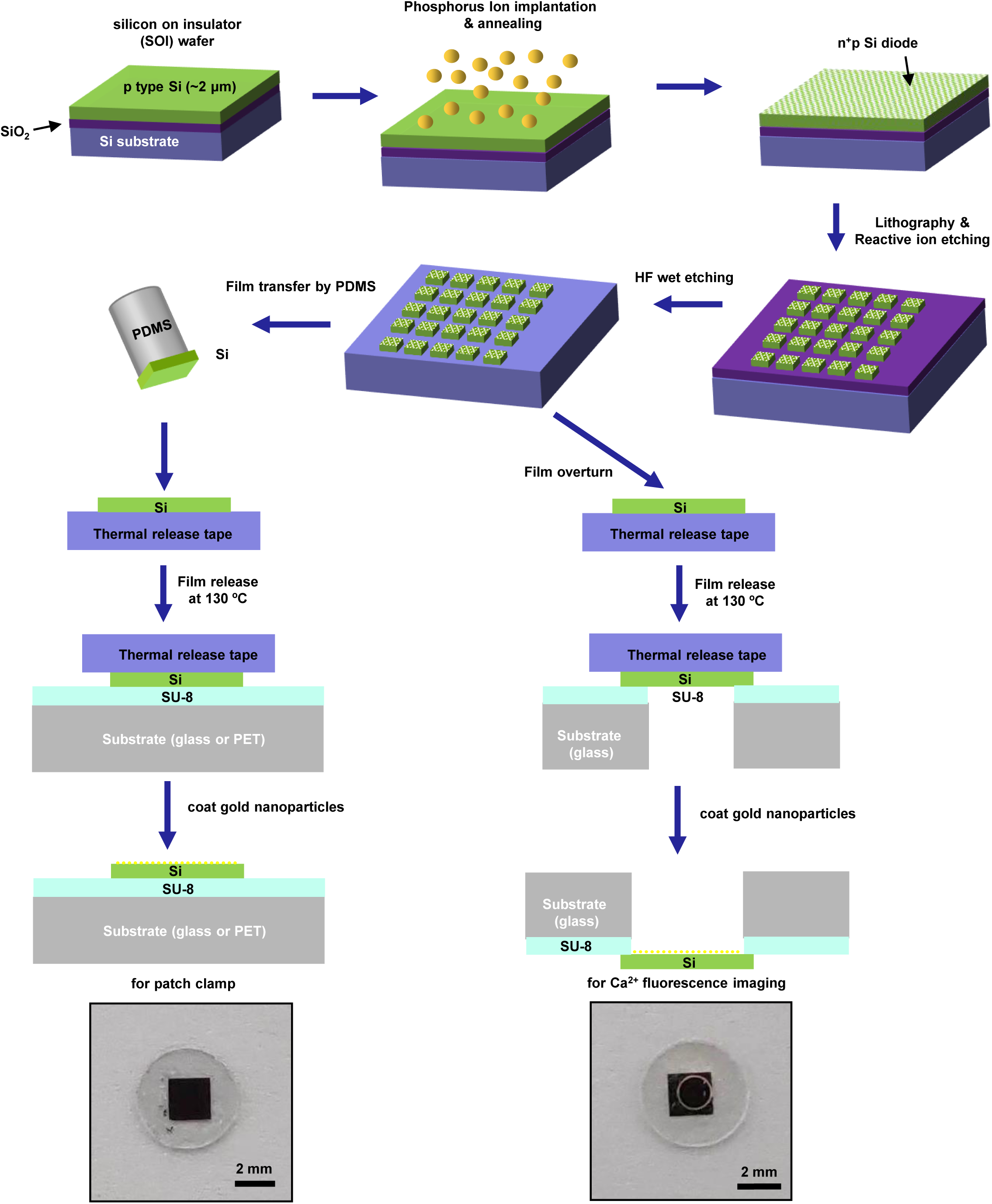
Processing flow for the fabrication of Au decorated n^+^p Si films used for (left) patch clamp and (right) for Ca^2+^ fluorescence imaging.

**Figure S2.**
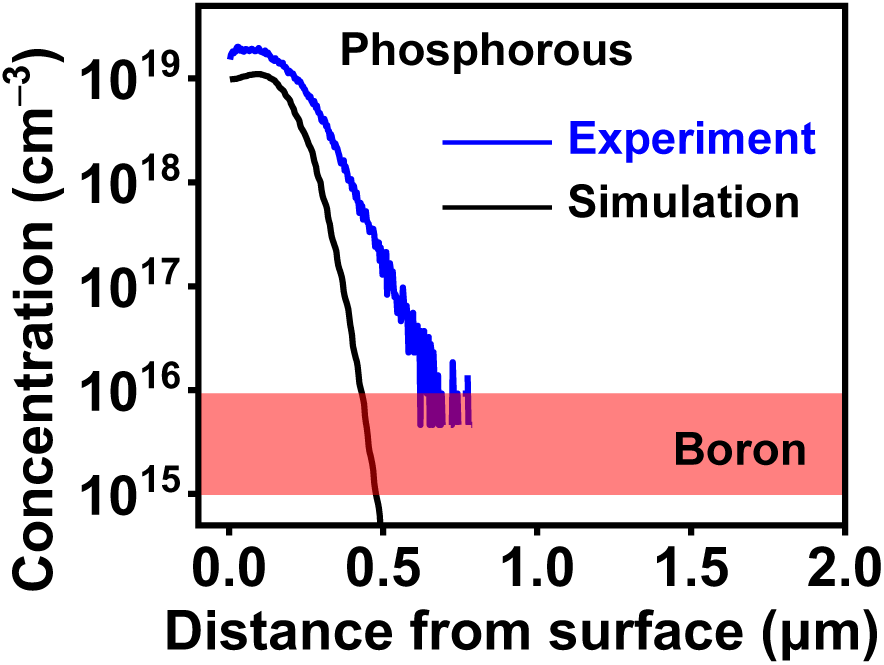
Doping profiles of the fabricated n^+^p Si film. Blue and black lines indicate phosphorous (P) concentration as a function of depth by secondary ion mass spectrometry (SIMS) measurement and Monte-Carlo simulation, respectively.

**Figure S3.**
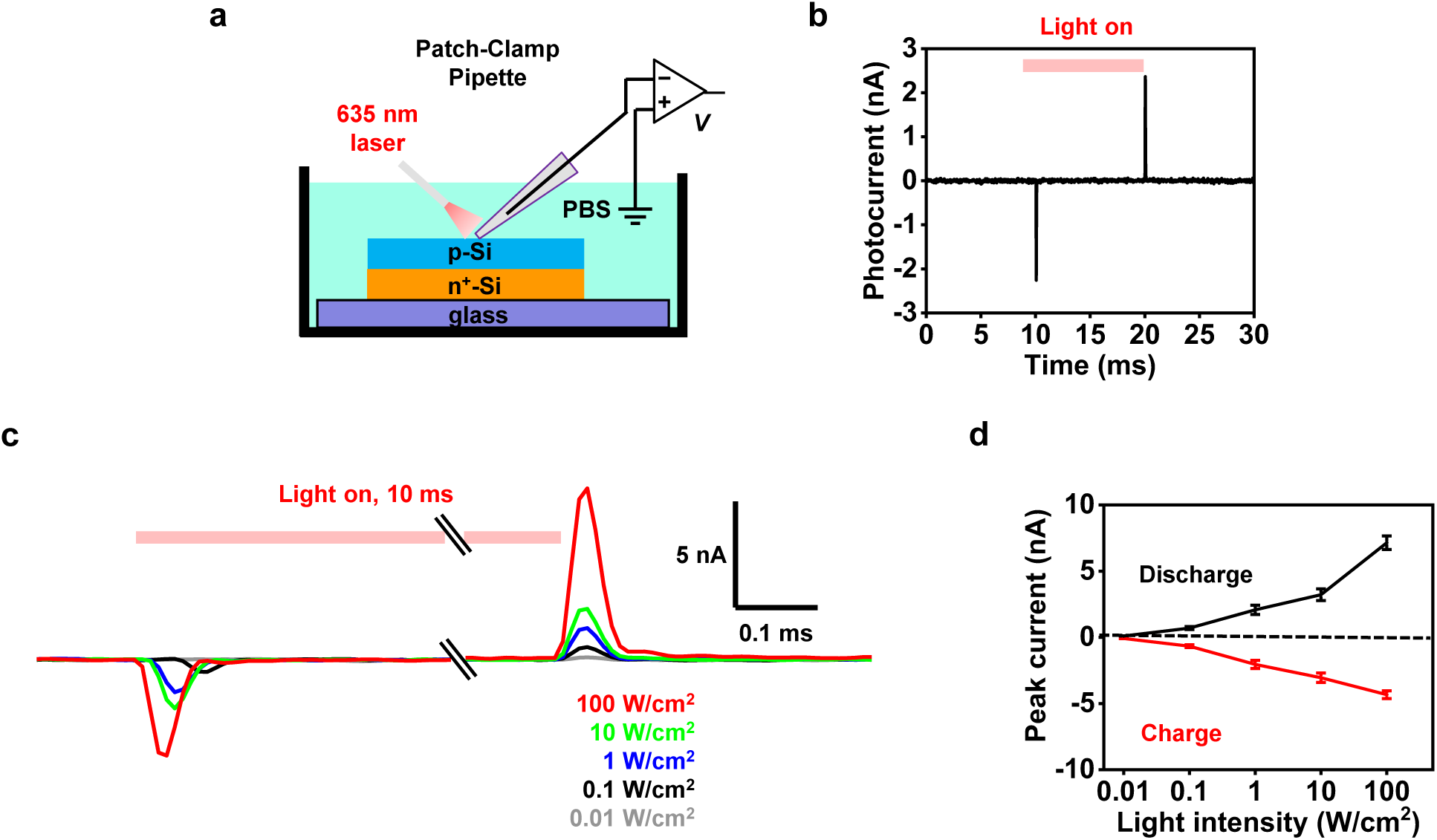
In vitro optoelectronic response of an n^+^p Si diode film without AuNPs. (a) (b) Scheme of the patch-clamp setup to measure the photocurrent of the Si film. (b) Photocurrent response for the Si film. The illumination condition is: intensity 10 W/cm^2^, pulse duration 10 ms. (c) Exploded view of charge and discharge peak currents for the Si film under illumination with different intensities (0.01, 0.1, 1, 10, 100 W/cm^2^). (f) Peak currents of charge and discharge as a function of light intensities. Data are presented as mean ± s.e.m..

**Figure S4.**
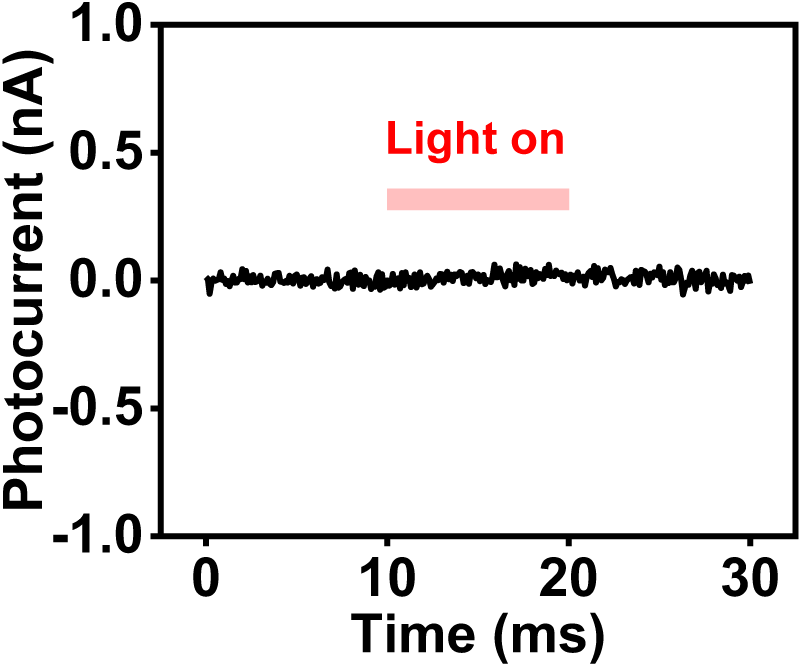
Optoelectronic response of a glass substrate. The illumination conditions is: intensity 10 W/cm^2^, pulse width 10 ms.

**Figure S5.**
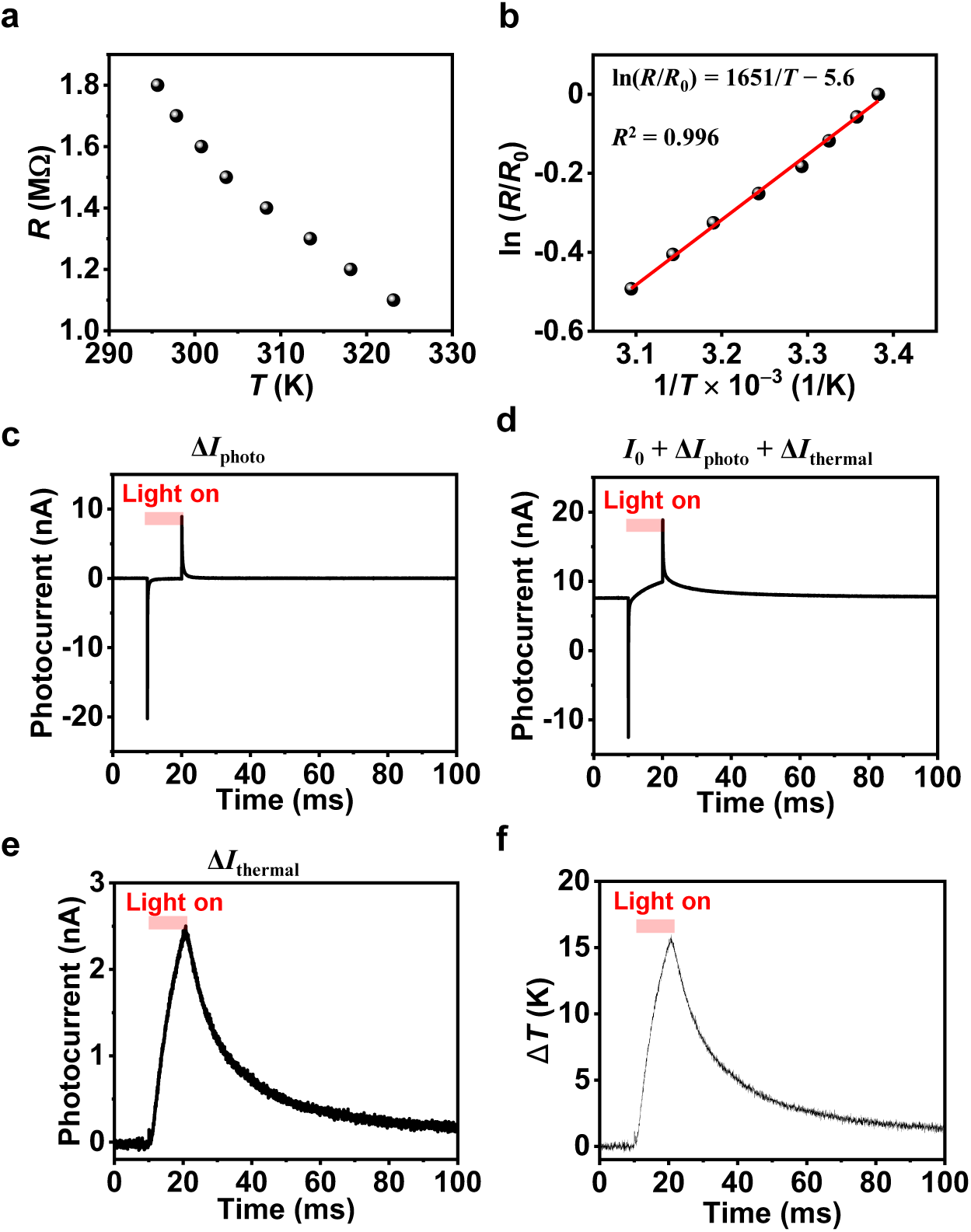
Characterization of photothermal effects for the Au decorated n^+^p Si diode film with the patch-clamp setup. (a) Measured resistance (*R*) of the pipette electrode versus temperature (*T*) in the PBS solution. (b) Arrhenius curve showing the linear relationship between ln(*R*/*R*_0_) and 1/*T*. *R*_0_ is the resistance of micropipette at room temperature (295 K). Black dots denote the measured values and the red line denotes the linear fit. (c) Transient photocurrent response (Δ*I*_photo_) without bias current (*I*_0_). The illumination condition is: pulse duration 10 ms and intensity 100 W/cm^2^. (d) Transient photocurrent response (*I*_0_ + Δ*I*_photo_ + Δ*I*_thermal_) with a bias current (*I*_0_ = 7.5 nA). (e) Calculated photothermal current (Δ*I*_thermal_) by subtracting data in (c) from data in (d). (f) Calculated dynamic temperature increase based on the calibration curve in (b). The temperature resolution is ∼0.5 K.

**Figure S6.**
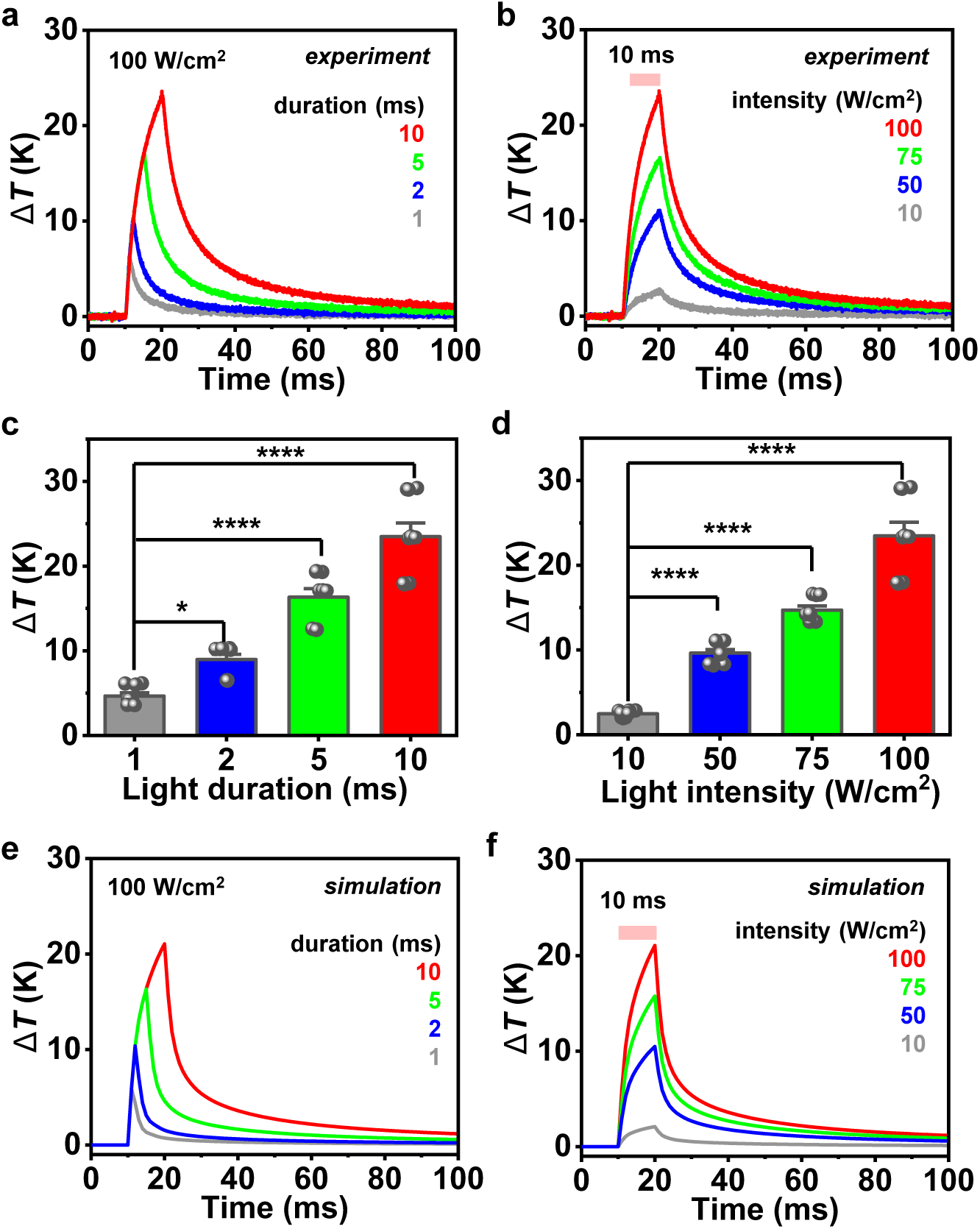
In vitro photothermal response of a Si diode film without AuNPs in PBS. (a, b) Measured transient temperature increase of the Si film illuminated by a pulsed laser at 635 nm, (a) with an intensity of 100 W/cm^2^ and different pulse durations (1, 2, 5, 10 ms), (b) with a pulse duration of 10 ms and different intensities (10, 50, 75, 100 W/cm^2^). (c, d) Statistical data of the maximum temperature rise related to conditions in (a) and (b) (*n* = 3 devices, 3 trials for each). (e, f) Simulated transient temperature response of a Si film corresponding to experimental conditions in (a) and (b). Data are presented as mean ± s.e.m and analyzed by one-way RM ANOVA. \**p* < 0.05, \*\**p* < 0.01, \*\*\**p* < 0.001, \*\*\*\**p* < 0.0001, n.s., not significant.

**Figure S7.**
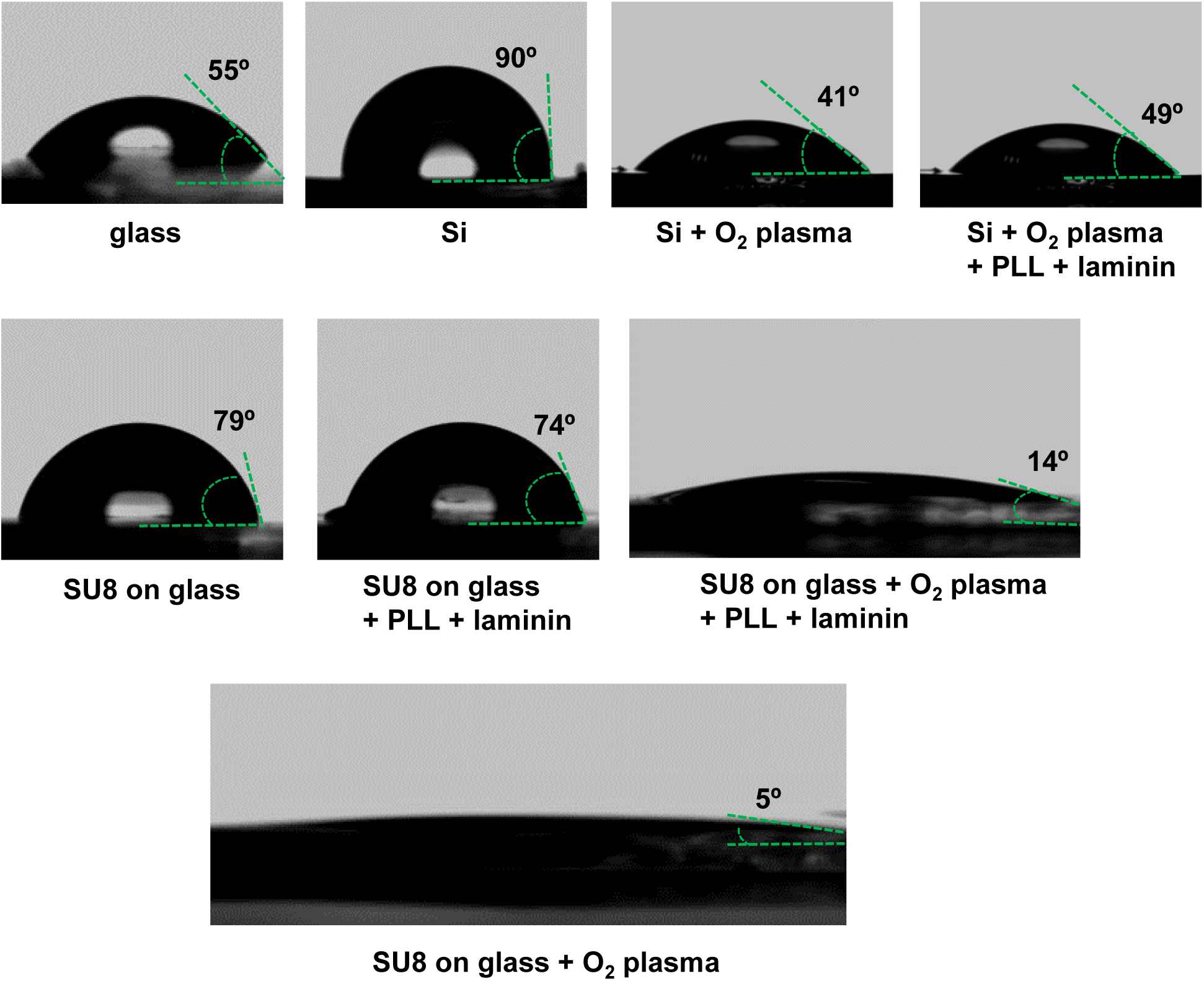
Photographs showing contact angles for water droplets (2 μl) on different substrates.

**Figure S8.**
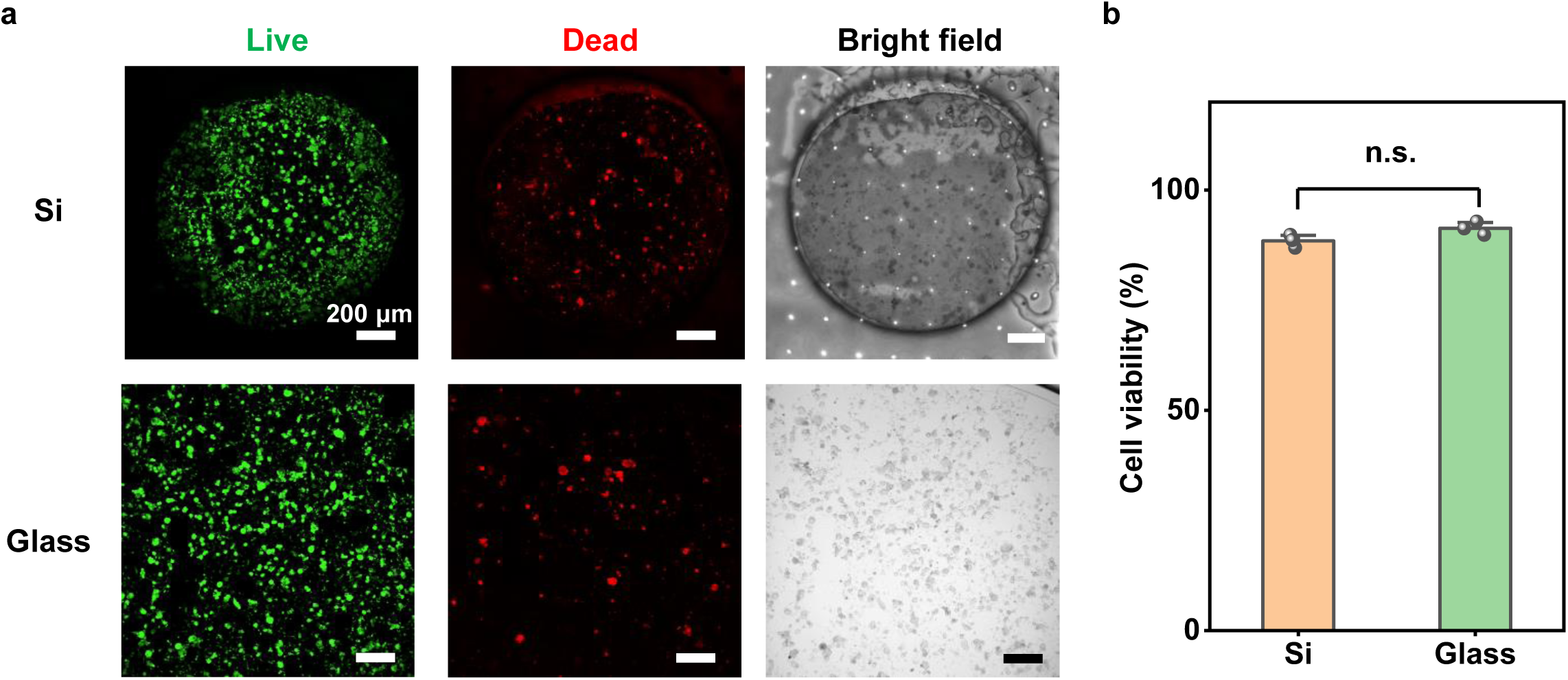
Biocompatibility of Si films. (a) Live/dead fluorescence assay performed on DRG neurons cultured on the Au coated n^+^p Si film and glass. Green (Calcien AM) and red (ethidium homodimer-1) denote live cells and dead cells, respectively. (b) Averaged viability (%) of DRG neurons cultured on the Si film and glass. Data are presented as mean ± s.e.m. (*n* = 3 images) and analyzed by unpaired *t*-test. n.s., not significant.

**Figure S9.**
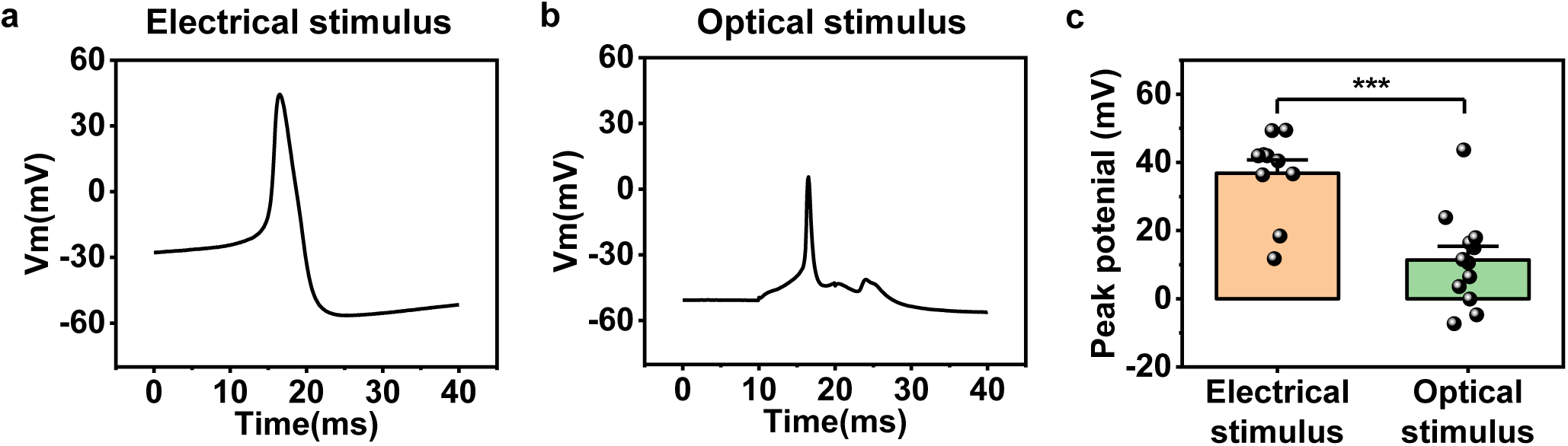
Comparision of typical APs initiated by electrical and optical stimuli. (a) A typical AP initiated by elevating the rest membrane potential to −30 mV. (b) A representative trace of AP activated by applying pulsed light (intensity 100 W/cm^2^, duration 10 ms, frequency 1 Hz) on the Si film. (c) Statistical results comparing the peak potentials of spikes by electrical and optical stimuli (electrical stimulus: *n* = 10 neurons, and optical stimulus: *n* = 12 neurons). Data are presented as mean ± s.e.m. and analyzed by unpaired *t*-test. \**p* < 0.05, \*\**p* < 0.01, \*\*\**p* < 0.001, \*\*\*\**p* < 0.0001, n.s., not significant.

**Figure S10.**
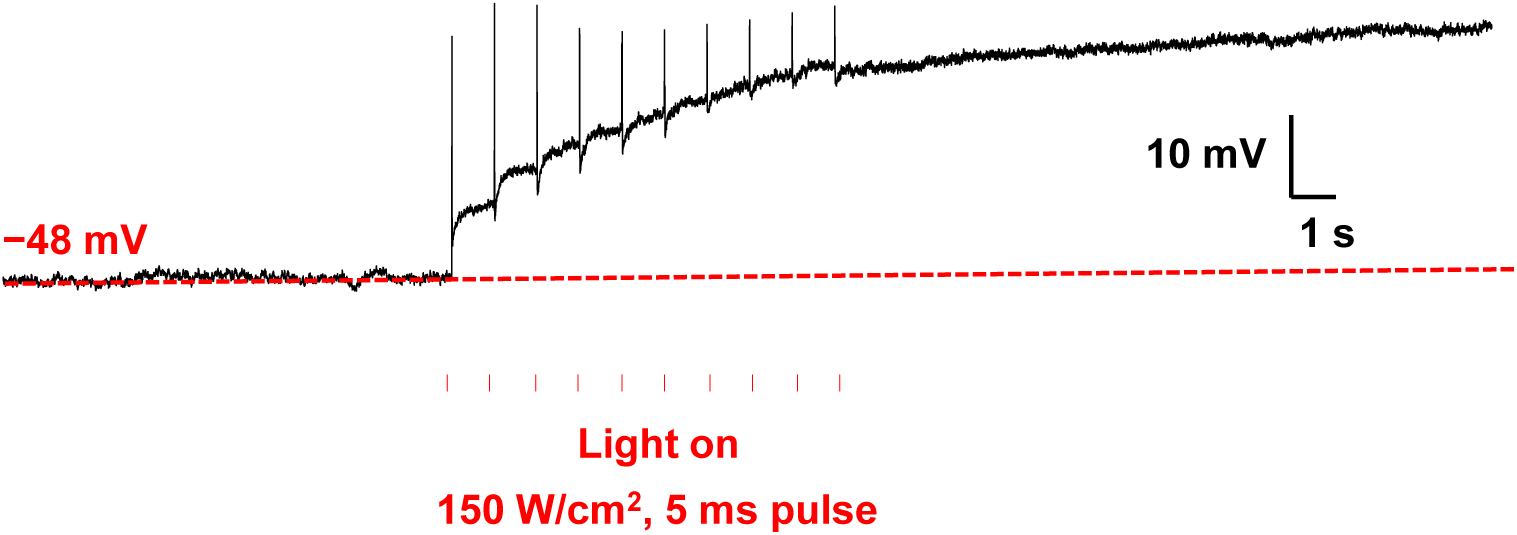
Recording of a DRG neuron on the Si film under high-power illumination. The illumination condition is: 635 nm laser, intensity 150 W/cm^2^, duration 5 ms, frequency 1 Hz. The strong phototherml effect eventually leads to the cell death.

**Figure S11.**
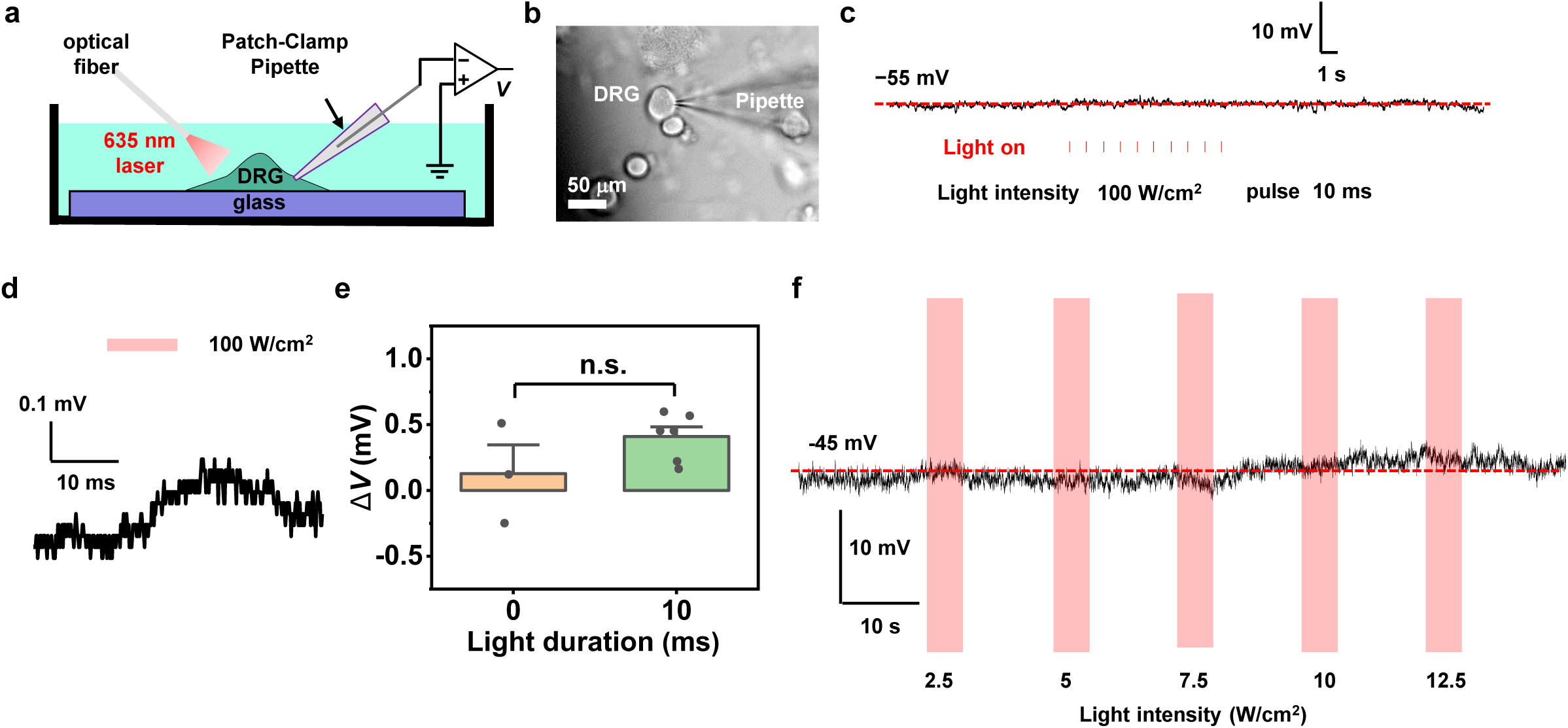
Electrophysiological activities of cultured DRG neurons on glass. (a) Schematic illustration of the patch-clamp setup to record electrophysiological signals of DRG neurons cultured on glass under light stimulation (with a 635 nm laser, ∼0.5 mm spot size). (b) Microscopic image of a DRG cell recorded by a pipette electrode. (c, d) Representative trace of membrane potentials under pulsed light: (c) intensity 100 W/cm^2^, duration 10 ms, frequency 1 Hz, (d) intensity 100 W/cm^2^, duration 10 ms. (e) Statistics of averaged membrane potentials in response to pulsed light stimulation (intensity 100 W/cm^2^) (*n* = 3 neurons, 2 trials for each). Data are presented as mean ± s.e.m. and analyzed by unpaired *t*-test. n.s., not significant. (f) Representative trace showing membrane potentials of the same DRG cell modulated by continuous illumination (duration 5 s, intensities 2.5, 5, 7.5, 10, 12.5 W/cm^2^).

**Movie S1.**
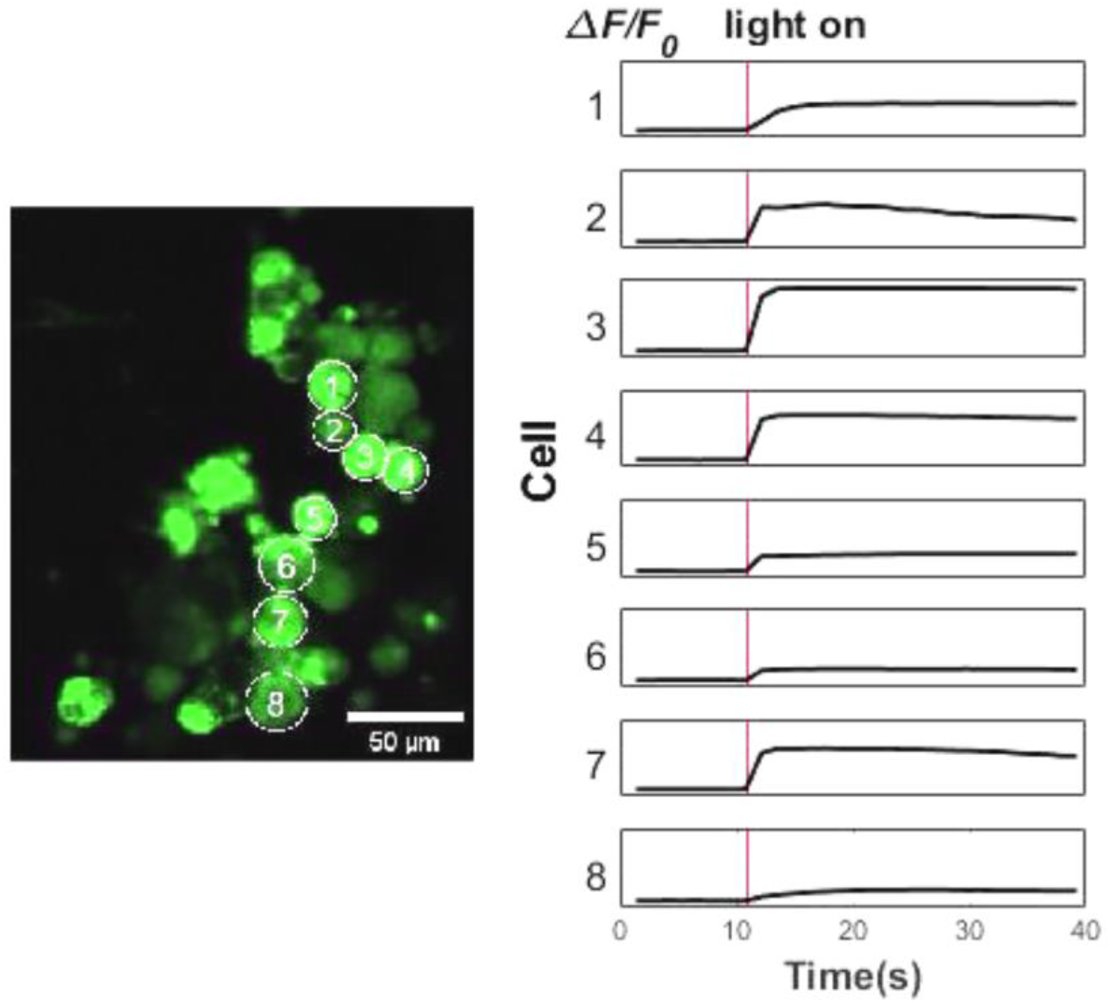
Video (5× speed) shows increased Ca^2+^ fluorescence of cultured DRG neurons evoked by a Si diode film under pulsed light stimulation (intensity 100 W/cm^2^, duration 10 ms).

**Movie S2.**
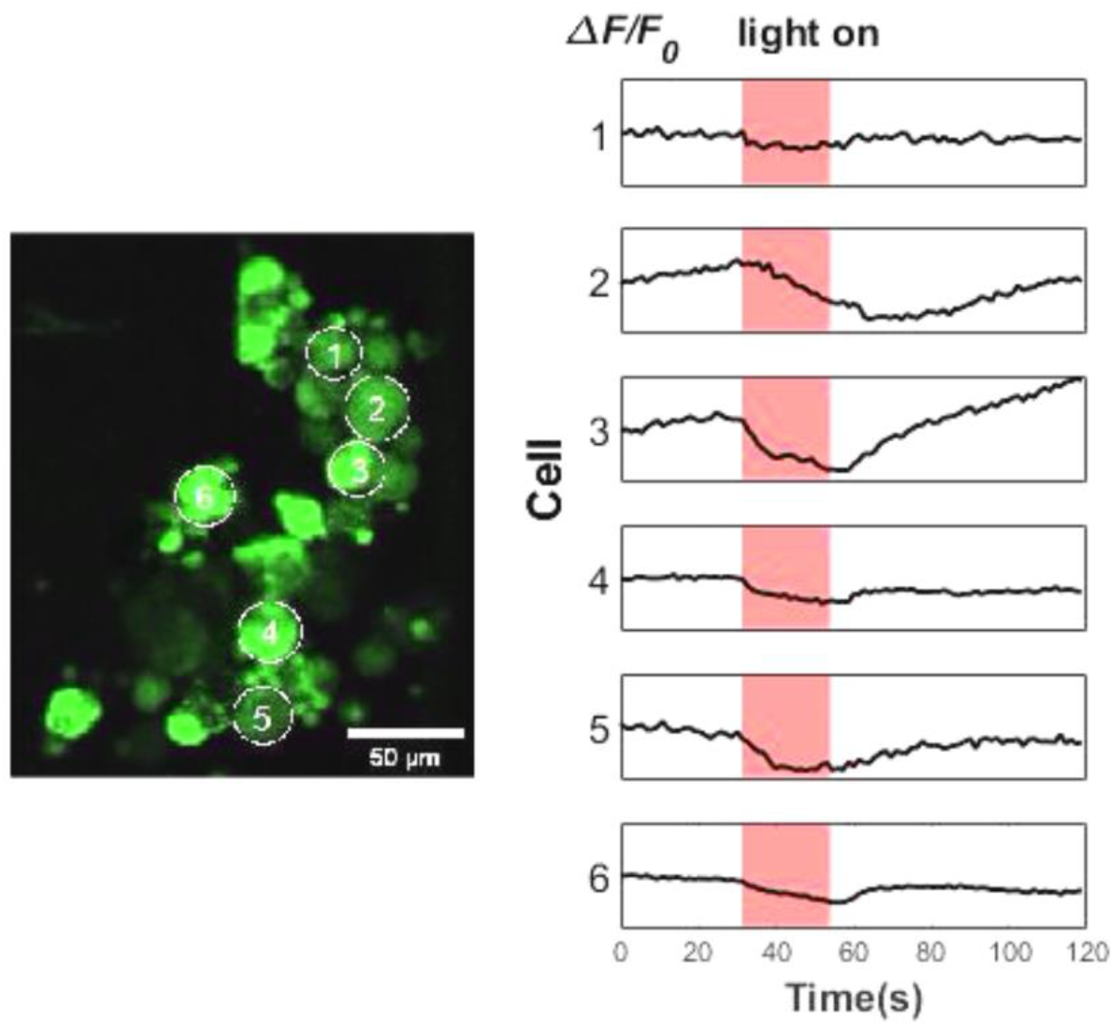
Video (10× speed) shows decreased Ca^2+^ fluorescence of cultured DRG neurons suppressed by the Si diode film under continuous light stimulation (intensity 1 W/cm^2^, duration 20 s).

## Author contributions

X.F. and X.S. developed the concepts. X.F., Y.H., G.T., B.Z., X.C., Y.W. and L.L. performed material design, fabrication and characterization. X.F., S.H. and J.L. performed patch clamp. X.F., W.L. and M.L. performed calcium imaging. Z.H. and L.M. performed in vivo electrophysiology. J.C. and H.W. performed simulations. J.M., S.S., L.Y., H.Z., X.L. and X.S. provided tools and supervised the research. X.F., Z.H. and X.S. wrote the paper in consultation with other authors.

## Data and materials availability

All data are available in the main text or the supplementary materials.

